# Identifying cellular markers of focal cortical dysplasia type II with cell-type deconvolution and single-cell signatures

**DOI:** 10.1101/2023.05.22.541770

**Authors:** Isabella C. Galvão, Ludmyla Kandratavicius, Lauana A. Messias, Maria C. P. Athié, Guilherme R. Assis-Mendonça, Marina K. M. Alvim, Enrico Ghizoni, Helder Tedeschi, Clarissa L. Yasuda, Fernando Cendes, André S. Vieira, Fabio Rogerio, Iscia Lopes-Cendes, Diogo F. T. Veiga

**Affiliations:** Department of Translational Medicine, School of Medical Sciences, University of Campinas, Campinas (UNICAMP), Brazil; Department of Pathology, School of Medical Sciences, University of Campinas (UNICAMP), Campinas, Brazil; Department of Neurology, School of Medical Sciences, University of Campinas (UNICAMP), Campinas, SP, Brazil; Department of Structural and Functional Biology, Institute of Biology, University of Campinas (UNICAMP), Campinas, Brazil; The Brazilian Institute of Neuroscience and Neurotechnology (BRAINN), Campinas, Brazil

**Keywords:** Focal cortical dysplasia, cell-type deconvolution, RNA-seq, astrogliosis

## Abstract

Focal cortical dysplasia (FCD) is a brain malformation that causes medically refractory epilepsy. FCD is classified into three categories based on structural and cellular abnormalities, with FCD type II being the most common and characterized by disrupted organization of the cortex and abnormal neuronal development. In this study, we employed cell-type deconvolution and single-cell signatures to analyze bulk RNA-seq from multiple transcriptomic studies, aiming to characterize the cellular composition of brain lesions in patients with FCD IIa and IIb subtypes. Our deconvolution analyses revealed specific cellular changes in FCD IIb, including neuronal loss and an increase in reactive astrocytes (astrogliosis) when compared to FCD IIa. Astrogliosis in FCD IIb was further supported by a gene signature analysis and histologically confirmed by glia fibrilla acidic protein (GAP) immunostaining. Overall, our findings demonstrate that FCD II subtypes exhibit differential neuronal and glial compositions, with astrogliosis emerging as a hallmark of FCD IIb. These observations, validated in independent patient cohorts and confirmed using immunohistochemistry, offer novel insights into the involvement of glial cells in FCD type II pathophysiology and may contribute to the development of targeted therapies for this condition.

## INTRODUCTION

Focal cortical dysplasia (FCD) is a malformation characterized by structural and cellular abnormalities of the cerebral cortex, such as disrupted organization of the neuronal layers and abnormal cell development. The condition is often difficult to treat with available anti-seizure medication and may require surgical resection of the affected brain tissue. FCD is a major cause of epilepsy in children, accounting for up to 50% of epilepsy surgeries in this age group^1^. FCDs are classified into three categories, each with its distinct histopathology and genetic alterations^2,3^. Among them, FCD type II is the most common^3^ and is distinguished from other types of FCDs due to loss of cortical organization in addition to the presence of dysmorphic neurons and balloon cells, which are mixed lineage immature cellular entities expressing both neuronal and glial proteins. At the histopathology level, FCD II can be further sub-classified into FCD IIa and IIb, presenting with dysmorphic neurons only (IIa) or with both dysmorphic neurons and balloon cells (IIb). Some patients with FCD II harbor somatic mutations in the mTOR pathway, mostly in the FCD IIb subtype^3^.

Cell type deconvolution methods applied to gene expression data have been used to estimate the proportions of different cell types in heterogeneous tissues in both healthy and disease settings^4,5^. To infer cell type abundances, these methods use regression on a reference expression matrix containing a set of marker genes constructed from reference expression datasets, such as bulk RNA-seq of individual cell types. More recent deconvolution approaches have been adapted to use single-cell RNA sequencing (scRNA-seq) reference signatures^6–8^, which might help to detect rarer cell types in mixture samples.

The brain is a complex organ composed of several cell types including neurons, glial cells (astrocytes, oligodendrocytes, ependyma, and microglia), and endothelial cells. Previous studies assessing cell-type deconvolution specifically in brain tissues found that cell-type estimates obtained through deconvolution highly correlated with those obtained through experimental approaches such as single-nuclei RNA-seq^9^ and immunohistochemistry^10^. Thus, deconvolution has been shown to predict cell-type proportions in brain tissues accurately and can be useful for exploring the contribution of cell types in diseases such as FCD.

Due to their distinct histopathological and genetic features, FCD II subtypes are likely to have different cellular compositions. In this study, we applied cell-type deconvolution based on single-cell signatures to systematically characterize the heterogeneity of the cellular populations in tissue samples from patients with FCD IIa or IIb. We then applied immunochemistry techniques to validate our findings.

## MATERIALS AND METHODS

### RNA-seq data acquisition

FCD type II RNA-seq from Kobow et al^11^ was retrieved from the European Nucleotide Archive (accession code SRP188422) and downloaded using the enaBrowserTools v. 1.1.0 (https://github.com/enasequence/enaBrowserTools). Raw fastq RNA-seq from Assis-Mendonça et al^12^ was obtained from the authors.

### RNA-seq data processing

Fastq files were processed using the RNA-seq nf-core pipeline v. 3.8.1^13,14^. Briefly, raw reads were trimmed using TrimGalore and aligned to the human reference genome hg38 using STAR^15^, followed by gene-level quantification using RSEM^16^ tool based on the Gencode v. 40 human reference transcriptome. Only samples containing at least 15 million mapped reads with an RSEM mapping rate of at least 80% were further selected for downstream analyses. After quality filtering, the first dataset included 11 samples from Kobow et al^11^ and 8 samples from Assis-Mendonça et al^12^. RNA-seq data was further normalized using DESeq2^17^ to remove batch effects. For the second dataset (Zimmer et al^18^), we used pre-processed RNA-seq of 32 patients obtained from the authors since raw fastq files could not be obtained from the original publication. The available clinical information for all samples is provided in Supplementary Table 1.

### Cell-type deconvolution using CIBERSORTx

The module “Impute cell fractions” of the CIBERSORTx tool was applied with the following parameters: relative mode, S-mode batch correction, and disabled quantile normalization. The website version (https://cibersortx.stanford.edu) was applied to analyze the single-cell signatures CA, VL, NG, and LK (details in Figure S1A). The Docker version of CIBERSORTx was applied to analyze the MS reference signature due to the size limit of the reference signature files in the online version (1GB).

### Methylome-based deconvolution using EpiSCORE

We applied the EpiSCORE tool^19^ with default parameters to a set of FCD methylomes profiled using Methyl-seq^11^ using a brain-specific DNAm signature containing the promoter-level methylation of 113 cell-type-specific marker genes^20^. The mixture input matrix was created by obtaining the average gene-level methylation from Methyl-seq peaks located at the proximal promoter (1kb upstream) of the transcription start site.

### FCD tissue collection

We collected specimens from 8 patients with epilepsy who underwent surgery for intractable seizures at the Epilepsy Surgery Program at Hospital de Clínicas, University of Campinas. All of the patients had a neuropathological diagnosis of FCD IIa (n = 4) or FCD IIb (n = 4), according to the guidelines of the International League Against Epilepsy^3^. All procedures were approved by the University of Campinas’s Research Ethics Board, and written informed consent was obtained from all participants.

### Immunohistochemical analyses

Brain specimens were transversely cut oriented to the pia mater. Then, the tissues were fixed in buffered formalin (Sigma, St Louis, MO, USA). After 48-96 hours, specimens were paraffin-embedded for immunohistochemistry (IHC). IHC was performed with primary antibodies against the glial fibrillary acidic protein (GFAP), an astrocytic marker (1:100, clone 6F2, Dako/Agilent, cat#M0761, Santa Clara, CA, USA), and MAP2, a protein expressed by neurons (1:1000, clone M13, Thermo Fisher, cat#13-1500, Waltham, MA, USA). Antibodies specificity was verified and IHC was performed as described in Mota et al^21^ and Assis-Mendonça et al^12^. Briefly, paraffin-embedded sections (4 um) were incubated with each antibody (overnight; at room temperature). Afterward, a solution with the secondary antibody and peroxidase (AdvanceTMHRP®, Dako, cat#K4068, Glostrup, Denmark; or EnvisionTM Flex+, Dako, cat#K8002, Glostrup, Denmark) was added for 30 min at 37°C. 3,3-diaminobenzidine was used as a chromogenic substrate and the sections were counterstained with hematoxylin. Control sections were concurrently submitted to the same protocol except for the primary antibody.

### Quantitative analysis of IHC

For quantitative analysis of immunoreactivity, a CS2 Aperio ScanScope scanner (Aperio Technologies, Vista, CA, USA) was used to obtain digitized images. For each individual, ten representative fields (20x digital magnification) of the cortex immunoreacted for each marker (GFAP and MAP2) were evaluated. All digitized images were obtained under the same luminance and were further analyzed with ImageJ software, following the same criteria: (i) deconvolution of each image (separation of hematoxylin and DAB channels) was performed by using the IHC Profiler plugin^22^; (ii) the DAB channel was used for quantification with the threshold tool; (iii) for each image, a threshold value was established so that the histological findings were preserved in the most accurate way considering the original photo, that is, to exclude the low-intensity gray value of background staining and to allow the quantification of the positive-stained area detected in the soma and branches^23,12^. Groups were compared using the Wilcox rank-sum test in R.

## RESULTS

### Cell-type deconvolution uncovers cellular alterations in FCD IIa and IIb lesions

To estimate cell type proportions in patients with FCD IIa and IIb, we obtained bulk RNA-seq data from three independent transcriptomic studies and applied CIBERSORTx to obtain estimates of the brain-specific populations (Figure 1A). The first dataset included 10 FCD IIa and 9 FCD IIb samples, obtained after combining data provided by Kobow et al^11^ and Assis-Mendonça et al^12^. Before applying CIBERSORTx, RNA-seq was uniformly re-processed to obtain a gene expression matrix (see Methods), and normalized with DESeq2^17^ to remove batch effects between studies. The second dataset served as an independent validation cohort and included normalized gene-level expression data of 11 FCD IIa and 21 FCD IIb patients profiled by RNA-seq, obtained from Zimmer et al^18^. We used four single-cell reference signatures - named CA, VL, NG, and LK after the original studies - to perform deconvolution (Figure S1A). These signatures were obtained from Sutton et al^9^ and derived from single-nuclei RNA-seq data obtained from the human cerebral cortex. The single-cell signatures contained gene expression profiles of major brain cell types, including neurons, astrocytes, oligodendrocytes, microglia, oligodendrocyte progenitors, and endothelial cells. The CA signature showed the best fit in both patient groups, as demonstrated by the Pearson correlation between the actual RNA-seq gene expression and the reconstructed gene expression by CIBERSORTx (Figure S1B).

**Figure 1.**
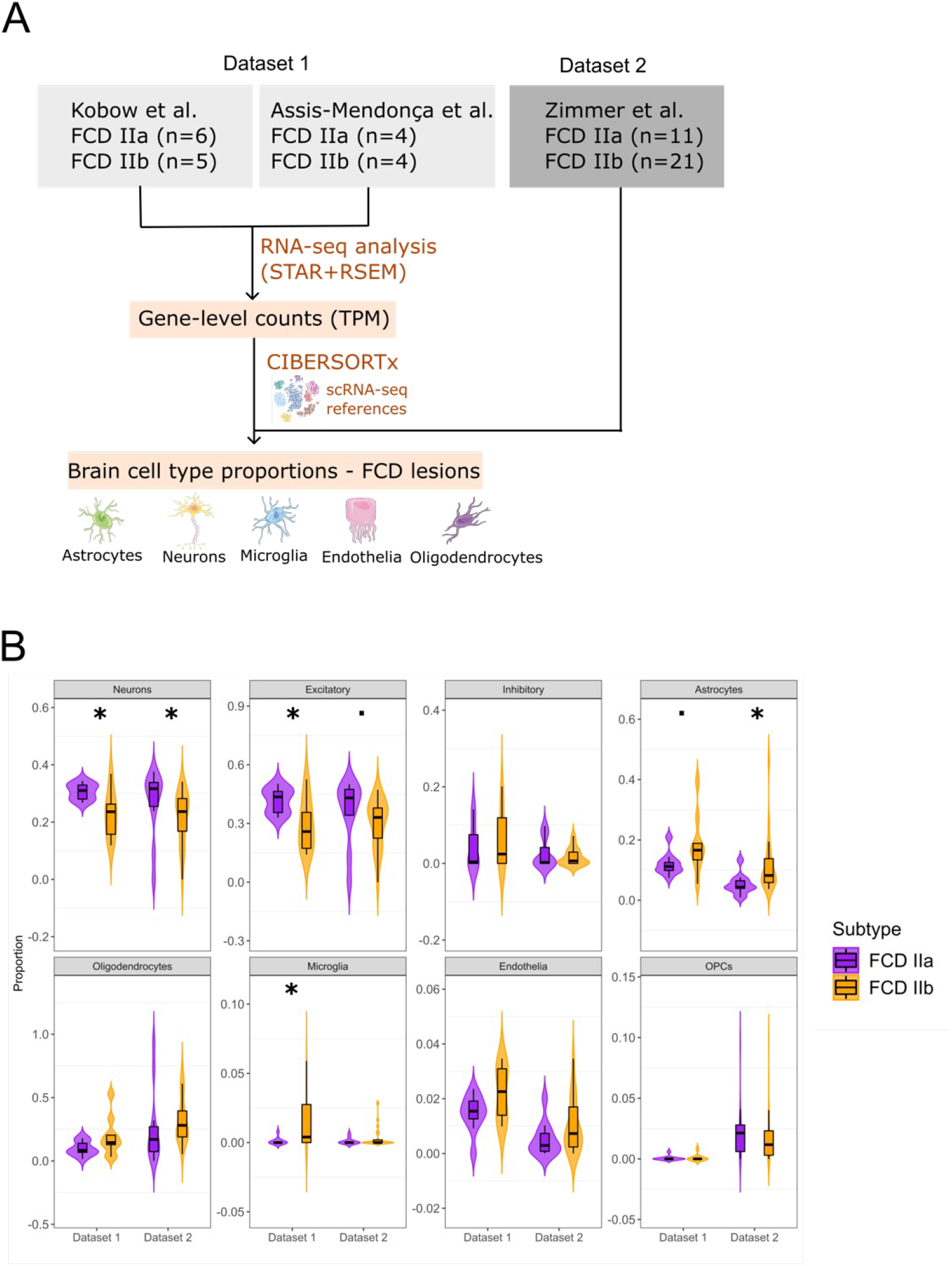
Cell-type deconvolution of major brain populations in FCD IIa and IIb lesions. **(A)** Schematic depiction of the analytical pipeline used for cellular deconvolution. RNA-seq datasets were obtained from three independent studies and were grouped into Datasets 1 and 2 (the number of samples is indicated). After RNA-seq data processing (see Methods), CIBERSORTx was performed to estimate cell type abundance using multiple single-cell reference signatures derived from snRNA-seq datasets of the human brain (see Fig. S1A). The Figure was partly generated using Servier Medical Art, provided by Servier, licensed under a Creative Commons Attribution 3.0 unported license. **(B)** Violin plot of cell type proportions in FCD IIa and IIb lesions in patients from Dataset 1 and Dataset 2, using the CA reference signature. The *y*-axis indicates the absolute cell type proportion (0 to 1) estimated by CIBERSORTx, while the *x*-axis indicates the patient dataset. The width of the violin indicates sample density, with the top, middle, and bottom of the black boxplot marking the 75^th^, 50^th^, and 25^th^ percentiles, respectively. Significant changes between cell type estimates in FCD IIa and IIb lesions were detected using the Wilcoxon rank-sum test. •p < 0,1, *p < 0.5, **p < 0.01, ***p < 10^−3^, ****p < 10^−4^.

In the first patient cohort (Dataset 1), CIBERSORTx found a significant decrease in neurons, and expansion of astrocytes and microglia populations in FCD IIb lesions when compared to FCD IIa (Figure 1B). The neuronal decrease was specific to the excitatory subtype, with no changes to inhibitory neurons (Figure 1B). The analysis of the independent patient cohort (Dataset 2) confirmed the estimated loss of excitatory neurons and higher astrocyte abundance in FCD IIb (Figure 1B). In addition, the decrease of excitatory neurons and increase of astrocytes was also detected by CIBERSORTx with other single-cell signatures VL, NG, and LK (Figures S1C-E).

Next, we wanted to determine which subtypes of excitatory neurons are affected in FCD IIb lesions. To address this question, we used CIBERSORTx and a single-cell signature containing 18 neuronal and non-neuronal cell types, annotated from an snRNA-seq dataset that profiled 48,920 individual nuclei from multiple sclerosis lesions and adjacent healthy tissue resected from the cortex^24^. The MS signature expanded upon the previous analysis by classifying neurons in different cortical layers into 6 excitatory and 5 inhibitory subtypes, as well as including a variety of glial and immune cells. The analysis found a specific decrease in excitatory neurons in the upper cortical layers 2/3 (cohort 1) and pyramidal subtypes (both patient cohorts), as shown in Figure 2A. Additionally, the deconvolution with the MS single-cell signature confirmed an increase in the astrocyte compartment in FCD IIb lesions in both patient groups (Figure 2B), consistent with the previous analyses using other signatures (Figures 1B, S1C-E).

**Figure 2.**
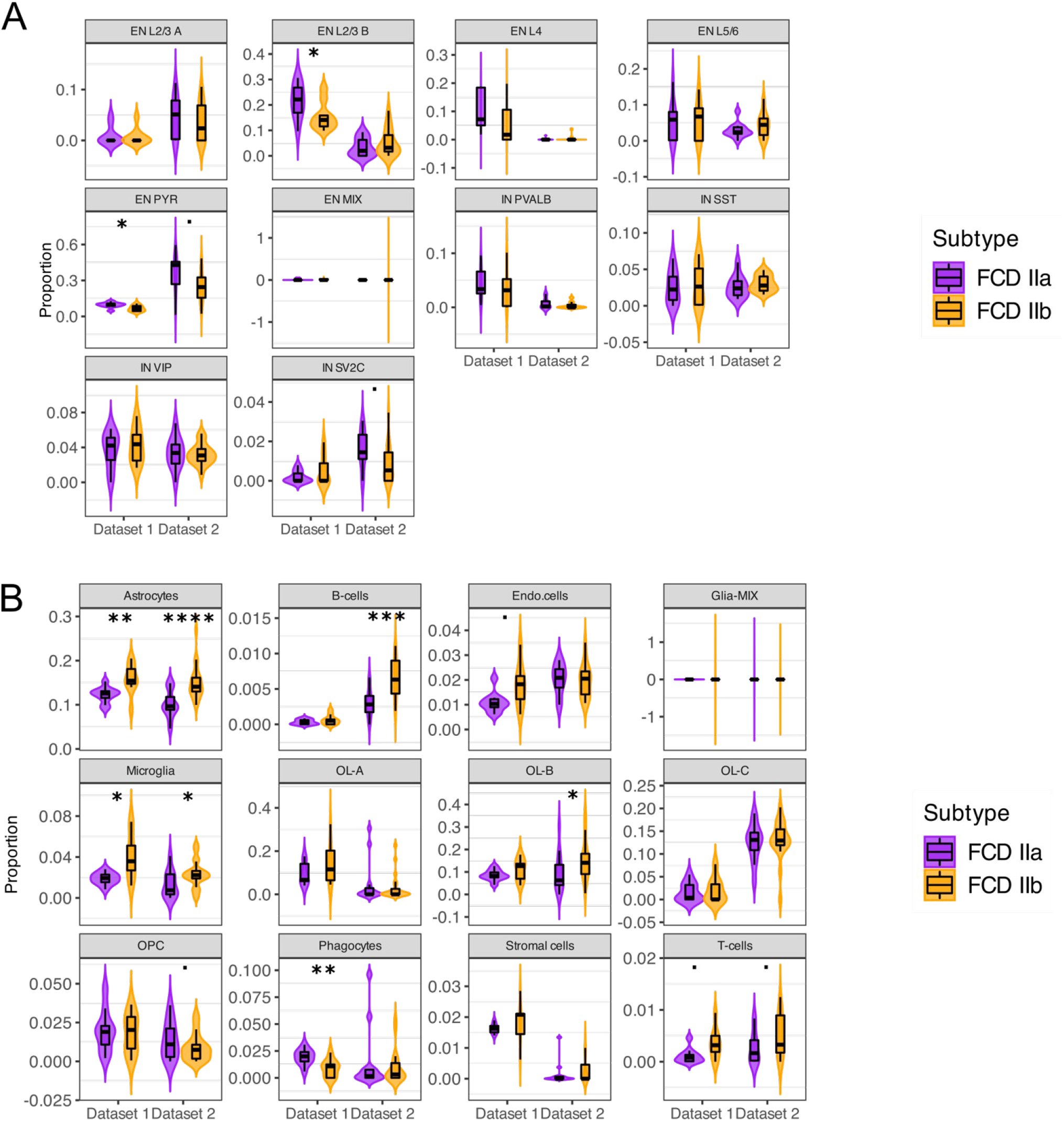
Cellular deconvolution of neuronal and non-neuronal subtypes in FCD IIa and IIb. **(A-B)** Violin plots of estimated cell type proportions in FCD IIa and IIb lesions of **(A)** excitatory and inhibitory neurons, and **(B)** non-neuronal cells. The *y*-axis indicates the absolute proportion (0 to 1) estimated by CIBERSORTx, while the *x*-axis indicates the patient dataset. Significant changes between cell type estimates in FCD IIa and IIb lesions were detected using the Wilcoxon rank-sum test. •p < 0,1, *p < 0.5, **p < 0.01, ***p < 10^−3^, ****p < 10^−4^. EN = excitatory neurons, IN = inhibitory neurons, and PYR = pyramidal neurons. L indicates the cortical layer (e.g. L2/3 refers to layers 2-3), Endo = endothelia, OL = oligodendrocytes, OPC = oligodendrocytes progenitor cells, A/B/C = subclusters A, B, C.

### Cellular alterations in FCD IIa and IIb inferred by methylome-based deconvolution

We also performed methylome-based deconvolution on a dataset of 6 FCD IIa and 5 FCD IIb samples profiled by Mehtyl-seq^11^ using the EpiSCORE tool^19^. This tool uses differentially methylated promoters to identify a set of cell-type marker genes (differentially expressed and specific to the cell type) in order to define tissue-specific signatures. Using this approach, Zhu et al^20^ proposed and validated brain-specific DNA methylation (DNAm) signature containing the promoter-level methylation of 113 cell type-specific marker genes present in brain cell types.

We found that unsupervised principal component analysis (PCA) using DNAm can distinguish the FCD IIa and IIb subtypes, indicating their distinct methylation profiles (Figure S2A). In addition, EpiSCORE estimated a decrease in neuron proportions and an increase in astrocytes in FCD IIb when compared to FCD IIa lesions (Figure S2B). Thus, these results were consistent with the findings from transcriptome-based deconvolution with CIBERSORTx.

### FCD IIb lesions are characterized by enhanced reactive astrogliosis

The above cell type deconvolution showed astrocyte expansion in FCD IIb. To further investigate the role of astrocytes in FCD, we evaluated astrogliosis in FCD II subtypes using a gene signature containing 18 biomarkers associated with astrocyte activation in response to deleterious stimuli associated with neurological conditions^25^. This astrogliosis gene signature included cytoskeleton genes GFAP, vimentin (VIM), and nestin (NES), which are often upregulated in reactive astrocytes. It also included genes involved in cell signaling, such as the transcription factor STAT3, the Ca^2+^ binding transporter S100B, and secreted proteins such as the C3 complement factor and SERPINA3.

Overall, we found that most astrogliosis biomarkers had increased expression in FCD IIb when compared to FCD IIa, and these genes were sufficient to group FCD II lesions by subtype (Figure 3A). Using the FGSEA enrichment test^26^, we confirmed that the astrogliosis gene signature is highly expressed in the FCD IIb subtype (Figure 3B). RNA-seq measurements of GFAP and VIM (vimentin) gene expression were significantly higher in FCD IIb in both patient cohorts, while NES (nestin) was higher in the first cohort (Figures 3C-D).

**Figure 3.**
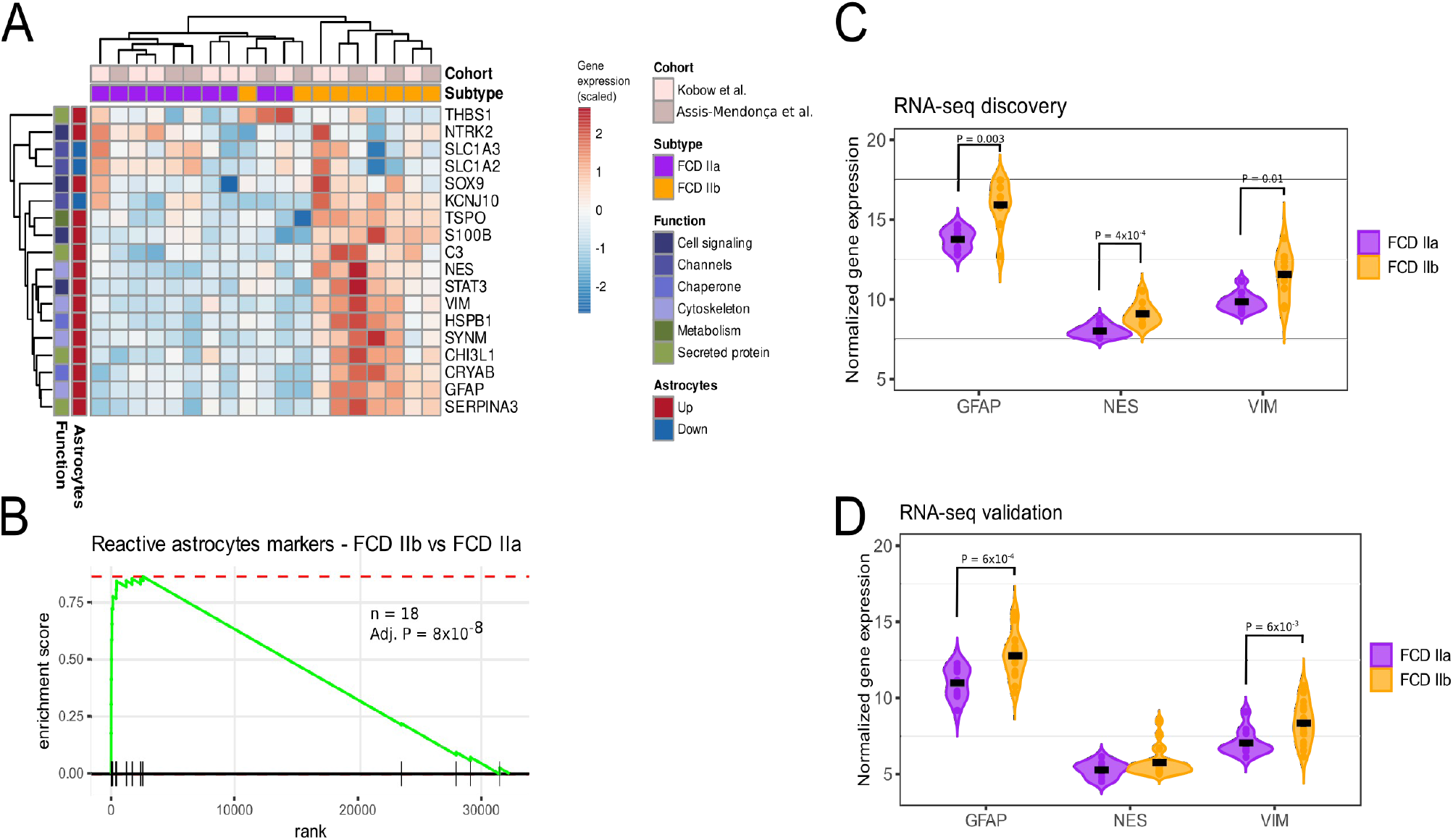
Reactive astrogliosis in FCD IIb lesions. **(A)** Heatmap expression of RA gene biomarkers in FCD IIb and IIa. Patient information (study and diagnosis) are indicated in the columns, and biomarker function and status (Up/Down-regulated in reactive astrocytes) are shown in the rows. **(B)** FGSEA plot of the RA signature (n = 18 genes, shown in A) based on expression FCD IIb. The *y*-axis indicates the enrichment score, and the *x*-axis represents all genes in the transcriptome ranked by log_2_ fold-change between FCD IIb and IIa. Genes in the RA signature are depicted by vertical lines along the *x*-axis. **(C-D)** Violin plots of GFAP, NES, and VIM gene expression in FCD IIa and IIb patients from RNA-seq data of **(C)** patient cohort 1 and **(D)** patient cohort 2. Significant changes between gene expression in FCD IIa and IIb were detected using the Wilcoxon rank-sum test (P-value indicated). Overlay dots represent gene expression in individual patients, and the black crossbar represents the median gene expression.

To confirm our findings, we performed IHC staining for the astrocyte marker GFAP and neuronal marker MAP2 on cortical tissues from 8 patients with intractable epilepsy and a diagnosis of FCD IIa (n=4) or IIb (n = 4, Figure 4). The results showed that GFAP immunostaining was significantly higher in FCD IIb compared to FCD IIa specimens (Figures 4A-G), indicating astrogliosis is more pronounced in FCD IIb lesions. In addition, MAP2 immunostaining levels were significantly lower in FCD IIb (Figures 4H-N), suggesting that astrogliosis is also accompanied by neuronal loss in FCD IIb.

**Figure 4.**
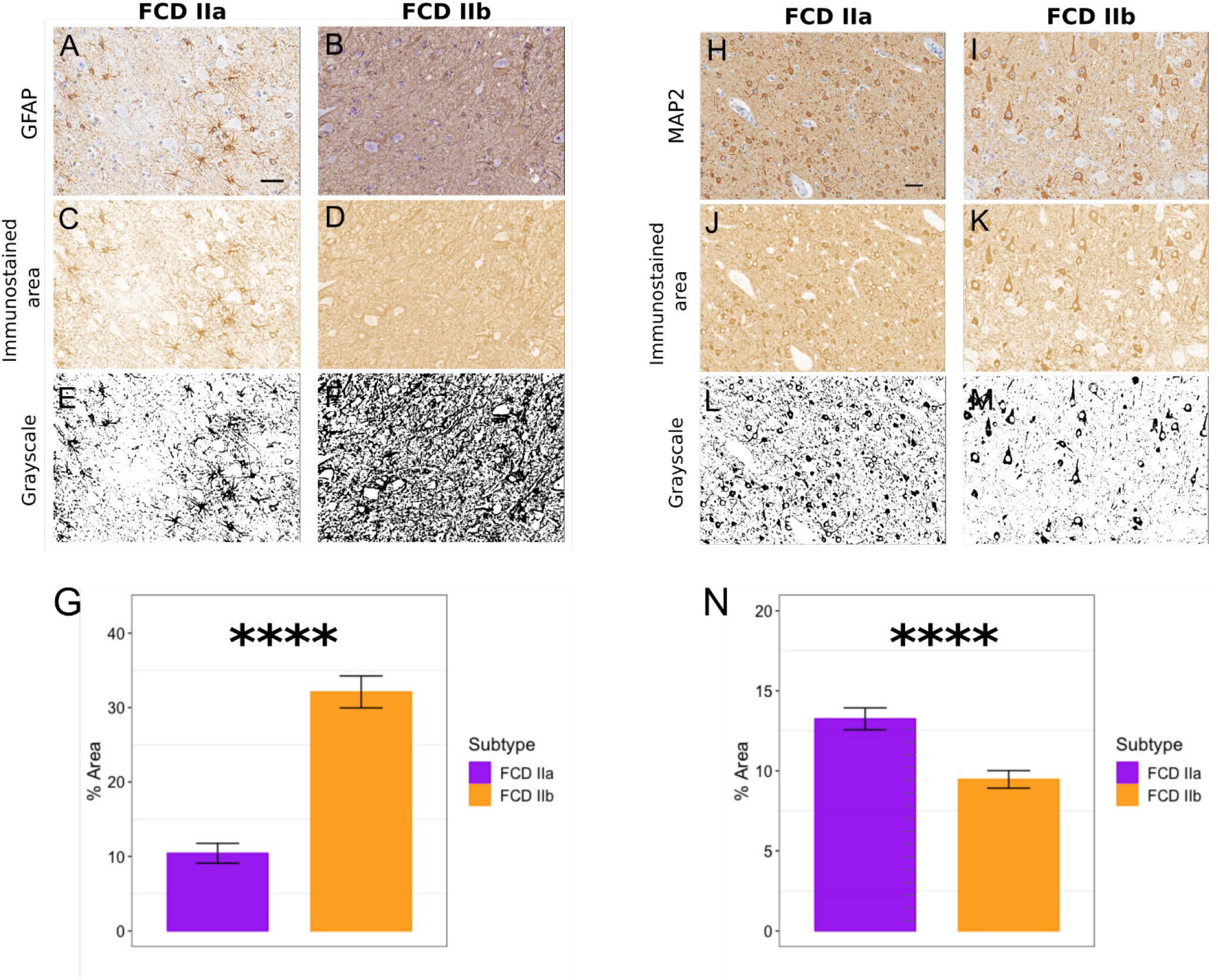
Immunostaining for GFAP and MAP2 proteins in FCD IIa and IIb tissues. Representative images of cortical immunostaining, DAB channel conversion, and grayscale conversion for **(A-F)** the astrocyte marker GFAP (glial fibrillary acidic protein), and **(H-K)** the neuron marker MAP2 (microtubule-associated protein 2). Barplots of GFAP **(G)** and MAP2 **(N)** expression levels in FCD IIa (n=4) and IIb (n=4) lesions. The mean expression was averaged from 10 randomly selected subfields in each lesion. The uncertainty bars represent 95% confidence intervals of the means. The image grayscale conversion of the positive immunostained area was used for marker quantification. Wilcoxon rank-sum test, ****p < 10^−4^. Scale bar: 50 μm.

## DISCUSSION

Cellular deconvolution is a powerful tool for analyzing transcriptomic and epigenetic data from complex tissues, where the contribution of different cell types to the overall gene expression profile is unknown or might be affected by disease. In this study, we leveraged this method to comprehensively characterize the previously unknown cell type composition of FCD type II, a highly epileptogenic condition that affects mainly children and adolescents.

Specifically, we applied gene expression deconvolution combined with single-cell signatures to estimate cell type proportions in patients diagnosed with FCD type IIa and IIb. We found that even though FCD type II lesions share common histological features, these FCD subtypes have remarkably distinct cellular profiles. Both transcriptome- and methylome-based cellular deconvolution analyses have shown that FCD IIb lesions are characterized by neuronal loss and astrocyte activation. In addition, our data showed that an astrogliosis gene signature is upregulated in FCD IIb. Further validation by IHC staining using MAP2 and GFAP markers confirmed these neuronal and glial changes in FCD IIb. Taken together, these findings indicate that neuronal loss and enhanced astrogliosis are hallmarks of FCD IIb lesions that might be included as criteria for subtype classification.

It is noteworthy that T2-weighted fluid-attenuated inversion recovery (FLAIR) magnetic resonance imaging of FCD IIb lesions reveals hyperintense signals in the cortical and subcortical regions, while FCD IIa lesions exhibit mild or absent hyperintense signals^27^. The discrepancy in FLAIR images between FCD II subtypes may be attributed to their cellular differences, even though further investigation is needed to explore this hypothesis.

Currently, the presence of balloon cells is the only histopathological feature used to distinguish between FCD type IIa and IIb lesions, and the status of glial cells is not taken into account in subtype classification. Based on our findings, we propose that measuring astrogliosis status by GFAP staining could be useful as an additional criterion for distinguishing between FCD IIb and FCD IIa. The analysis of GFAP status might be applied in conflicting cases where the clinical or neuroimaging findings suggest a type IIb lesion and the inspected tissue sample does not exhibit balloon cells^12^.

Our computational analysis was designed to be as robust and comprehensive as possible. We searched public repositories to obtain all available RNA-seq and methylome data for FCD type II for which clinical annotation was available. We utilized both discovery and validation patient cohorts, containing transcriptomic data generated in three research centers. Finally, to rule out the effect of the underlying single-cell signature in cellular deconvolution, we reproduced the results using four independent single-cell signatures derived from single-cell studies of the cerebral cortex, which is the main site of tissue changes in FCD.

In conclusion, our results indicate that the cellular environment in FCD IIb lesions is distinct from that of FCD IIa, with a higher level of astrogliosis in FCD IIb. This enhanced astrocyte response may contribute to increased neuroinflammation and the observed neuronal loss in FCD IIb. Further research is necessary to fully understand the role of astrocytes in FCD IIb and to investigate the potential therapeutic benefits of targeting astrocytes in the treatment of this condition.

## Data availability

RNA-seq analyzed in this study are available at the European Nucleotide Archive (accession code SRP188422) and at the Gene Expression Omnibus (accession code GSE213488) repositories. RNA-seq from Zimmer et al^18^ was obtained from the authors.

## Acknowledgments

D.F.T.V. was supported by a Young Investigator award from Fundação de Amparo à Pesquisa do Estado de São Paulo (FAPESP), Brazil (grant numbers 2019/07382-2 and 2020/15112-2). I.C.G was supported by a FAPESP fellowship (grant number 2022/01530-2). I.L.-C. was supported by Conselho Nacional de Pesquisa (CNPq), Brazil (grant number 311923/2019-4), and Coordenação de Aperfeiçoamento de Pessoal de Nível Superior (CAPES), Brazil (grant number 001). M.C.P. A. was supported by a fellowship from FAPESP (grant number 2020/04780-4). F.R. was supported by FAPESP (grant number 2019/08259-0). C.L.Y. was supported by Conselho Nacional de Pesquisa (CNPq), Brazil (grant number 315953/2021-7). F.C. was supported by the FAPESP Research Innovation and Dissemination Center (grant number 2013/07559-3).

## Author contributions

D.F.T.V., I.L-C., and F.R. contributed to the study design. I.C.G. and D.F.T.V. performed bioinformatic analyses. L.K. and F.R. performed immunohistochemistry experiments and data analysis. G.R.A-M., M.K.M.A., E.G., H.T., L.A.M., M.C.P.A., C.L.Y., A.S.V., and F.C. provided reagents and feedback. D.F.T.V., I.C.G., L.K., and F.R. drafted figures and the manuscript. All authors reviewed and approved the final version of the manuscript.

## Competing interests

The authors declare no competing interests.

## SUPPLEMENTARY FIGURES

**Figure S1.**
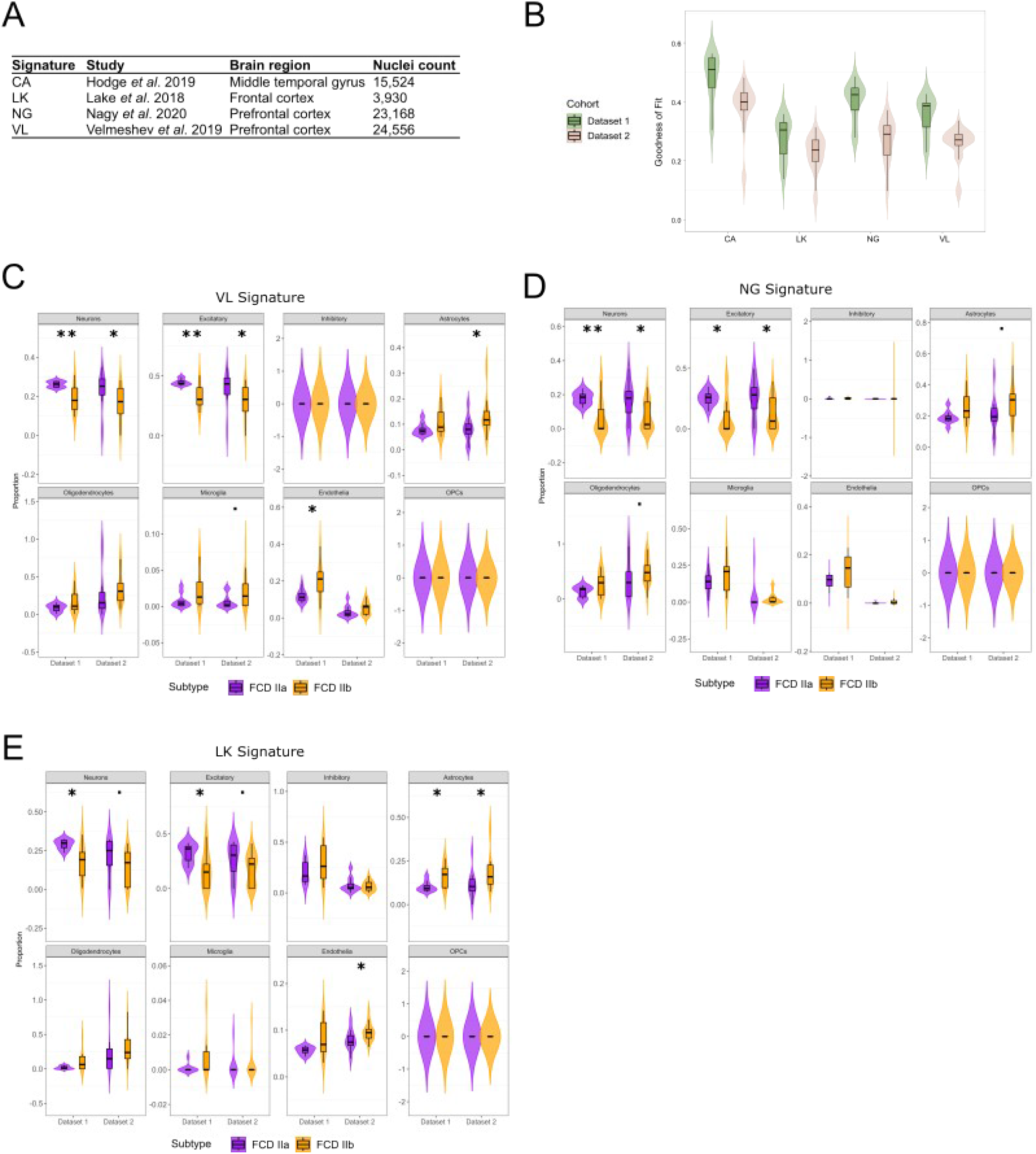
Cellular deconvolution of major brain cell types in FCD IIa and IIb using multiple single-cell reference signatures. **(A)** Single-cell reference signatures used for cell-type deconvolution, derived from single-nuclei RNA-seq studies from regions of the cerebral cortex. **(B)** Violin plots of the CIBERSORTx goodness-of-fit (i.e. Pearson correlation between the actual RNA-seq gene expression and reconstructed gene expression) for the reference signatures CA, VL, NG, and LK in Dataset 1 and Dataset 2. The width of the violin indicates sample density, with the top, middle, and bottom of the white boxplot marking the 75th, 50th, and 25th percentiles, respectively. **(C-E)** Violin plots of estimated cell type proportions in FCD IIa/IIb lesions in patients from Dataset 1 using the **(C)** VL signature, **(D)** NG signature and **(E)** LK signature. The *y*-axis indicates the absolute proportion (0 to 1) estimated by CIBERSORTx, while the *x*-axis indicates the patient dataset. The width of the violin indicates sample density, with the top, middle, and bottom of the white boxplot marking the 75th, 50th, and 25th percentiles, respectively. Significant changes between cell-type estimates in FCD IIa/IIb lesions were detected using the Wilcox rank-sum test. •P < 0,1, *P < 0.5, **P < 0.01, ***P < 10^−3^, **** P < 10^−4^.

**Figure S2.**
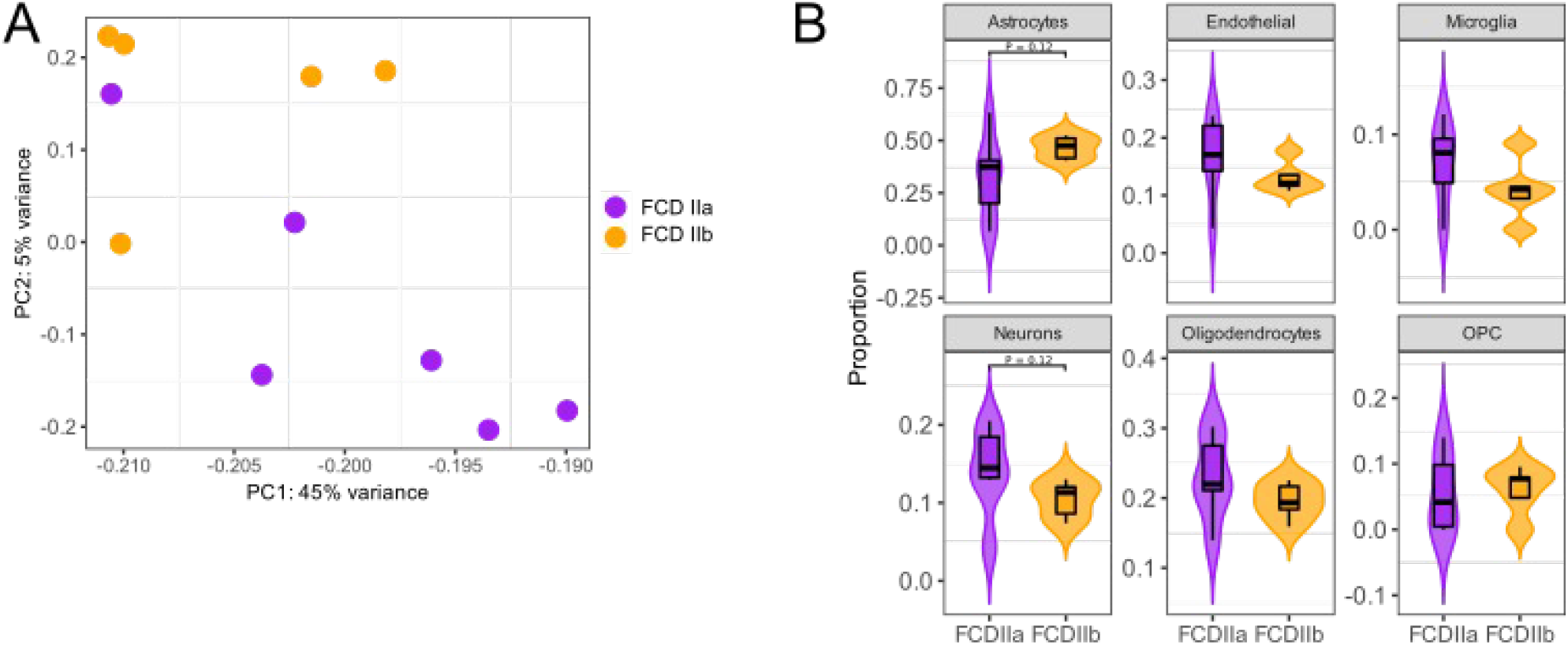
Cellular deconvolution based on FCD methylomes. **(A)** Dot plots representation of FCD IIa/IIb methylomes after dimensionality reduction with principal component analysis (PCA). The *x*-axis and *y*-axis refer to the first and second PCA components, respectively. **(B)** Violin plots of cell type proportions estimated by EpiScore in FCD IIa/IIb methylomes. The *y*-axis indicates the absolute proportion (0 to 1), while the *x*-axis indicates the disease subtype. The width of the violin indicates sample density, with the top, middle, and bottom of the white boxplot marking the 75th, 50th, and 25th percentiles, respectively.

## Notes

### Competing Interest Statement

The authors have declared no competing interest.

